# Genome Size Estimation and Full-length Transcriptome of *Sphingonotus tsinlingensis:* Genetic Background for the Drought-Adapted Grasshopper

**DOI:** 10.1101/2021.03.08.434347

**Authors:** Lu Zhao, Hang Wang, Le Wu, Kuo Sun, De-Long Guan, Sheng-Quan Xu

**Affiliations:** College of Life Sciences, Shaanxi Normal University, Xi’an 710119, China

**Keywords:** *Sphingonotus* Fieber, grasshoppers, PacBio isoform sequencing, gene functions, genetic background

## Abstract

*Sphingonotus* Fieber, 1852 (Orthoptera: Acrididae) is a species-rich grasshopper genus with ~146 species. All species of this genus prefer dry environments, such as: desert, steppe, sand, and stony benchland. This genomic study aimed to understand the evolution and ecology of these grasshopper species. Here, the genome size of *Sphingonotus tsinlingensis* was estimated using flow cytometry and the first high-quality full-length transcriptome of this species is presented, which may serve as a reference genetic resource for the drought-adapted grasshopper species of *Sphingonotus* Fieber. The genome size of *Sphingonotus tsinlingensis* was ~12.8 Gb. Based on the 146.98 Gb Pacbio isoform sequencing data, 221.47 Mb full-length transcripts were assembled. Among these transcripts, 88,693 non-redundant isoforms were identified with an average length of 2,497 bp and an N50 value of 2,726 bp, which was much longer than the formal grasshopper transcriptome assemblies. A total of 48,502 protein coding sequences were determined, and 37,569 were annotated in public gene function databases. A total of 36,488 simple tandem repeats, 12,765 long non-coding RNAs, and 414 transcription factors were also identified. According to gene functions, 70 heat shock proteins and 61 P450 genes that may correspond to drought adaptation of *S. tsinlingensis* were identified. The genome of *Sphingonotus tsinlingensis* is an ultra-large and complex genome. Full-length transcriptome sequencing is an ideal strategy for genomic research. This is the first full-length transcriptome of the genus *Sphingonotus.* The assembly parameters were better than all known grasshopper transcriptomes. This full-length transcriptome may be used to understand its genetic background and the evolution and ecology of grasshoppers.

## 1 Introduction

*Sphingonotus* Fieber, 1852 (Orthoptera: Acrididae) is a species-rich grasshopper genus with approximately 146 species (Cigliano et al., 2021). The most fascinating feature of these species is that they prefer a dry environment, such as: the desert, steppe, sand, and stony benchland (Husemann et al., 2014). These grasshoppers are distributed in the arid zone of the northern hemisphere, from North Africa to Eastern Asia. How they have adapted to drought environments and bloomed into species-richness has attracted many studies (Husemann et al., 2015; García-Navas et al., 2017). *Sphingonotus tsinlingensis* (Orthoptera: Acridoidea) is a Chinese endemic grasshopper that is distributed in the sand and stony benchland along the Northern Qinling mountains (Tse-Ming, 1963). This species belongs to the oriental lineage of the wide-spread and species-rich genus *Sphingonotus,* represents the adaptive status of *Sphingonotus spp.* in the East, and may serve as an ideal model to provide molecular evidence for studying the evolution and ecology of grasshoppers in arid environments (Husemann et al., 2014; Moussi et al., 2018). However, only a few molecular sequences have been determined for this species and there are no studies on the functional genes involved in its drought adaptation (Cui and Huang, 2012; Shah et al., 2019). To facilitate these evolutionary and ecological studies, adequate and sufficient sequences are needed to as a genetic background.

Due to the rapid progress in sequencing technologies, especially long-read sequencing, genomics has become basic data for evolutionary and ecological studies on non-model organisms (Morin et al., 2018). However, it is still very expensive to sequence a whole genome for non-model organisms with a large genome (Meyer et al., 2021). The genome size (GS) of grasshoppers (Orthoptera: Acrididae) ranges from 3.84 Gb *(Melanoplus differentialis)* (Yasuzumi et al., 1958) to 16.93 Gb *(Podisma pedestris}* (Westerman et al., 1987), and comprises extensive repetitive elements. These characteristics make the whole genome assembly of grasshopper genomes challenging, and sequencing costs are very expensive (Wang et al., 2014; Verlinden et al., 2020). These large complex genomes also increase the difficulty in sequencing the transcriptome, so deeper sequence coverage is required. Moreover, the GS of *S. tsinlingensis* needs to be estimated before sequencing may occur.

Here, a transcriptome of *S. tsinlingensis* is presented for further functional and ecological studies where the deep-coverage Pacbio isoform sequencing technique has been used (Camacho et al., 2015; Yuan et al., 2019). Besides protein coding genes, different types of genetic motifs, such as transcription factors (TFs), simple sequence repeats (SSRs), and long noncoding RNAs (lncRNAs) were identified and classified. The full-length transcriptome of *S. tsinlingensis* was compared to other grasshoppers to evaluate its quality and heat shock proteins and P450 genes were analyzed for further investigations on the drought adaptation of these grasshoppers.

## 2 Materials and Methods

### 2.1 Sample collection

Adults of *S. tsinlingensis* were collected from the natural populations at pebbled beach in Xi’an (34°02’05.2”N 108°33’04.3”E) on September 14, 2020. Some grasshoppers were kept alive in insect mesh cages until flow cytometry (FCM) was performed in the laboratory, while others were dissected, immediately immersed in liquid nitrogen, and then kept at −80 °C.

### 2.2 GS estimation using FCM

FCM was used to investigate the GS of *S. tsinlingensis* (Dolezel and Bartos, 2005). In FCM, DNA content is in direct proportion to fluorescence intensity. That is, if the fluorescence intensity of an internal standard and target specimens is obtained, and the internal standard’s GS is known, the target specimens’ GS may be calculated. Here, *Locusta migratoria* (1C = 6.5 G) (Wang et al., 2014) has been chosen as the internal standard. The FCM protocol followed the methods of Gregory and Johnston (Gregory and Johnston, 2008) and Hare and Johnston (Hare and Johnston, 2011). To prepare single cell suspensions, head tissues of *L. migratoria* and *S. tsinlingensis* were removed and placed in a tissue grinder with 1 mL Galbraith buffer (Galbraith et al., 1983), and ground 20 times. Then, the solution was filtered through a 38 μm nylon mesh to remove cellular debris and stained with 50 μg/mL propidium iodide. The above steps were conducted on ice. Finally, the solution was stored at 4 ? for 30 mins in the dark. The GS was measured using a flow cytometer (Beckman Coulter Cytoflex S, Krefeld, Germany) with three technical replicates, set to activate with a 488 nm laser and low flow rate.

### 2.3 RNA extraction

Total RNA was separated from different tissues and extracted using the TRIzol reagent (Invitrogen, Carlsbad, CA, USA), following the manufacturer’s instructions. RNA degradation and contamination were evaluated using 1% agarose gels. The integrity and purity of RNA was determined using an Agilent 2100 Bioanalyzer (Agilent Technologies, CA, USA) and a NanoDrop 2000 (Thermo Scientific, Wilmington, DE, USA). Only total RNA samples with a RNA integrity number value ≥8 were used for the preparation and construction of PacBio or HiSeq sequencing libraries.

### 2.4 Library preparation and sequencing

The Isoform Sequencing library was prepared according to the Isoform Sequencing protocol using the Clontech SMARTer PCR cDNA Synthesis Kit and the BluePippin Size Selection System protocol as described by Pacific Biosciences (PN 100-092-800-03).

### 2.5 Data analysis

Sequence data were processed using the SMRT Link 5.1 software. Circular consensus sequences (CCSs) were generated from subread BAM files with the following parameters: min_length 50, max_drop_fraction 0.8, no_polish TRUE, min_zscore −9999.0, min_passes 2, min_predicted_accuracy 0.8, and max_length 15,000. CCS.BAM files were generated, which were then classified into full-length and non-full-length reads using pbclassify.py, ignorepolyA false, and minSeqLength 200. Non-full-length and full-length FASTA files were produced and fed into the cluster step, which performed isoform-level clustering, followed by final Arrow polishing, hq_quiver_min_accuracy 0.99, bin_by_primer false, bin_size_kb 1, qv_trim_5p 100, and qv_trim_3p 30. Additional nucleotide errors in consensus reads were corrected using the Illumina RNA sequencing data with the software LoRDEC (Salmela and Rivals, 2014). Any redundancy in corrected consensus reads was removed using the Cluster Database at High Identity with Tolerance (Fu et al., 2012) (-c 0.95 -T 6 -G 0 -aL 0.00 -aS 0.99) to obtain final transcripts for subsequent analysis (**Supplementary Figure S1**).

### 2.6 Gene functional annotation

Gene function was annotated based on the following databases: non-redundant protein sequences (NR) (Li et al., 2002), non-redundant nucleotide sequences (NT), protein family (Pfam), clusters of orthologous groups of proteins (KOG) (Tatusov et al., 2003), Swiss-Prot (Bairoch and Apweiler, 2000), Kyoto Encyclopedia of Genes and Genomes (KEGG) (Kanehisa et al., 2004), and gene ontology (GO) (Ashburner et al., 2000). The basic local alignment search tool (BLAST) was used with the e-value set to ‘1e-10’ in NT database analysis, whereas the Diamond BLASTX software was used with the e-value set to ‘1e-10’ in the NR, KOG, Swiss-Prot, and KEGG database analyses. The Hmmscan software was used in Pfam database analysis (**Supplementary Table S2)**.

### 2.7 Protein-coding sequence (CDS) prediction

The ANGEL (Shimizu et al., 2006) pipeline, a long-read implementation of ANGLE, was used to determine CDSs from cDNAs. This species, or closely related species, confident protein sequences were used for ANGEL training and then the ANGEL prediction was run for given sequences.

### 2.8 TF analysis

Animal TFs were predicted using the animalTFDB 2.0 database.

### 2.9 LncRNA analysis

The Coding-Non-Coding Index (Sun et al., 2013), Coding Potential Calculator (Kong et al., 2007), Pfam-scan (Finn et al., 2016) and ***p***redictor of ***l***ong non-coding RNAs and m***e***ssenger RNAs based on an improved ***k***-mer scheme (Li et al., 2014) tools were used to predict the coding potential of transcripts. Transcripts predicted with coding potential by either/all of the four tools above were filtered out, and those without coding potential were the candidate set of lncRNAs.

### 2.10 SSR analysis

SSRs of the transcriptome were identified using MIcroSAtellite (MISA) (v1.0; https://webblast.ipk-gatersleben.de/misa/), which allows the identification and localization of perfect microsatellites, as well as compound microsatellites that are interrupted by a certain number of bases.

## 3 Results

### 3.1 GS estimation of *S. tsinlingensis*

The GS of *S. tsinlingensis* was estimated using FCM. Two peaks were identified, in which P1 and P2 represented the counted cell intensities of *L. migratoria* and *S. tsinlingensis,* respectively. The mean fluorescent intensity of peak P1 and P2 was 8.11 × 10^5^ and 15.98 × 10^5^ (low coefficient of variation, less than 5%), respectively. The ratio of GS of *S. tsinlingensis* to *L. migratoria* was equal to that of P2 to P1. The GS of *S. tsinlingensis* was calculated to be approximately 12.81 Gb (**Figure 1**). This GS was much larger than *L. migratoria*, and was also a large one in the Acrididae.

**FIGURE 1.**
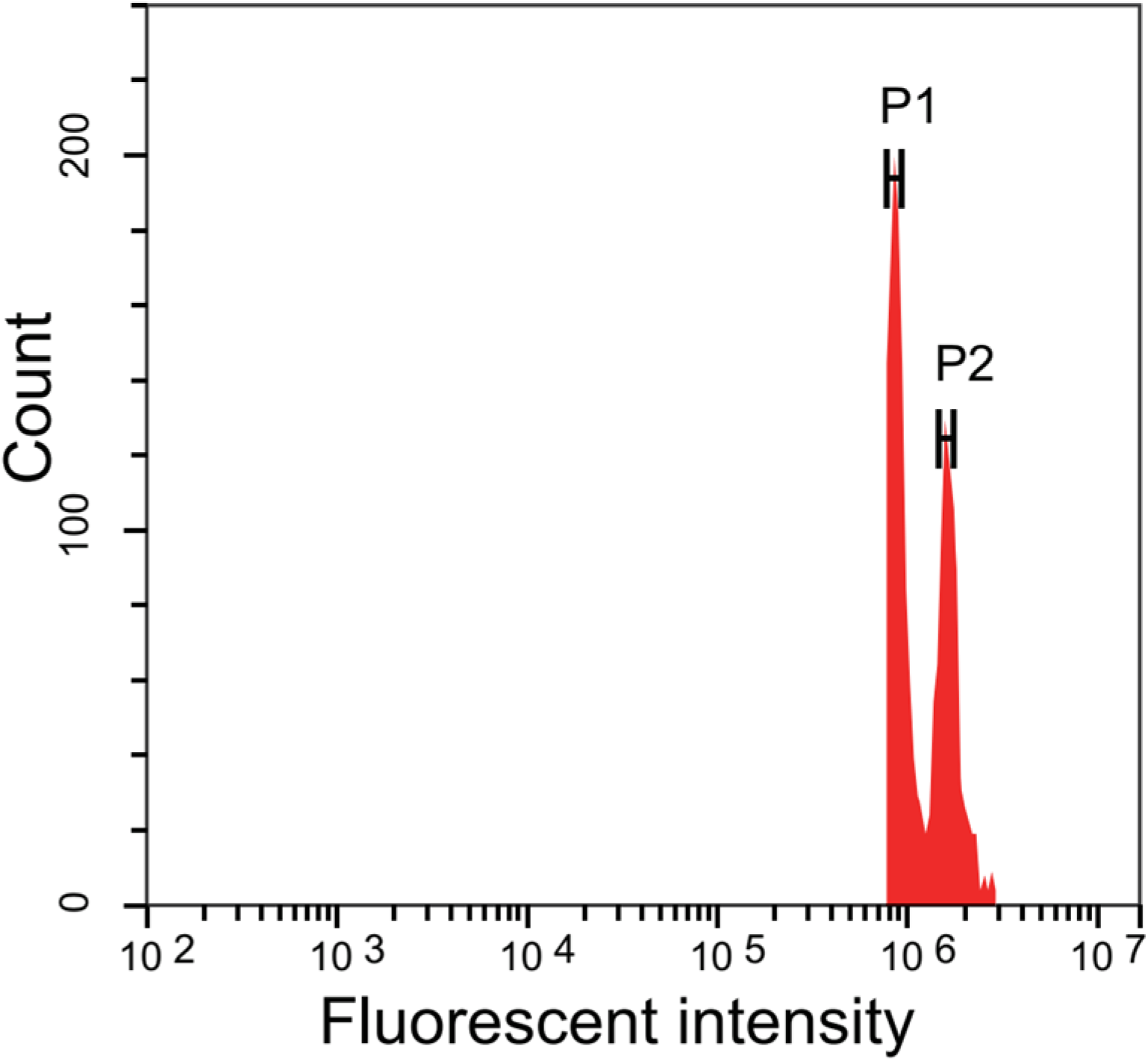
The genome size estimation of *S. tsinlingensis* using flow cytometry analysis. *Locusta migratoria* is the internal standard. P1 and P2 represent the peaks of *Locusta migratoria* and *Sphingonotus tsinlingensis,* respectively. The X-axis represents fluorescent intensity and the Y-axis represents counts of cells.

### 3.2 Full-length transcriptome assembly of *S. tsinlingensis*

A total of 146.98 Gb raw polymerase reads were obtained using the PacBio isoform sequencing platform. After filtering and self-correcting the raw data, 28.6 million subreads were processed into 901,383 CCS reads. Then, these CCSs were clustered and polished into 88,693 non-redundant fulllength, non-chimeric isoforms, which was the final molecular sequence pool for screening gene components (**Supplementary Table S1**). Compared to other reported grasshopper genomes and transcriptomes, the full-length transcriptome of *S. tsinlingensis* had better parameters. The identified number, average length, N50 value, and the percentage of long-non-redundant isoforms (over 1000 bp) were 88693, 2497 bp, 2726 bp and 99.28%, respectively. The most outstanding parameter for these results was the average transcript length, which was hundreds of bps more than the other transcriptomes (**Table 1**).

**TABLE 1.**
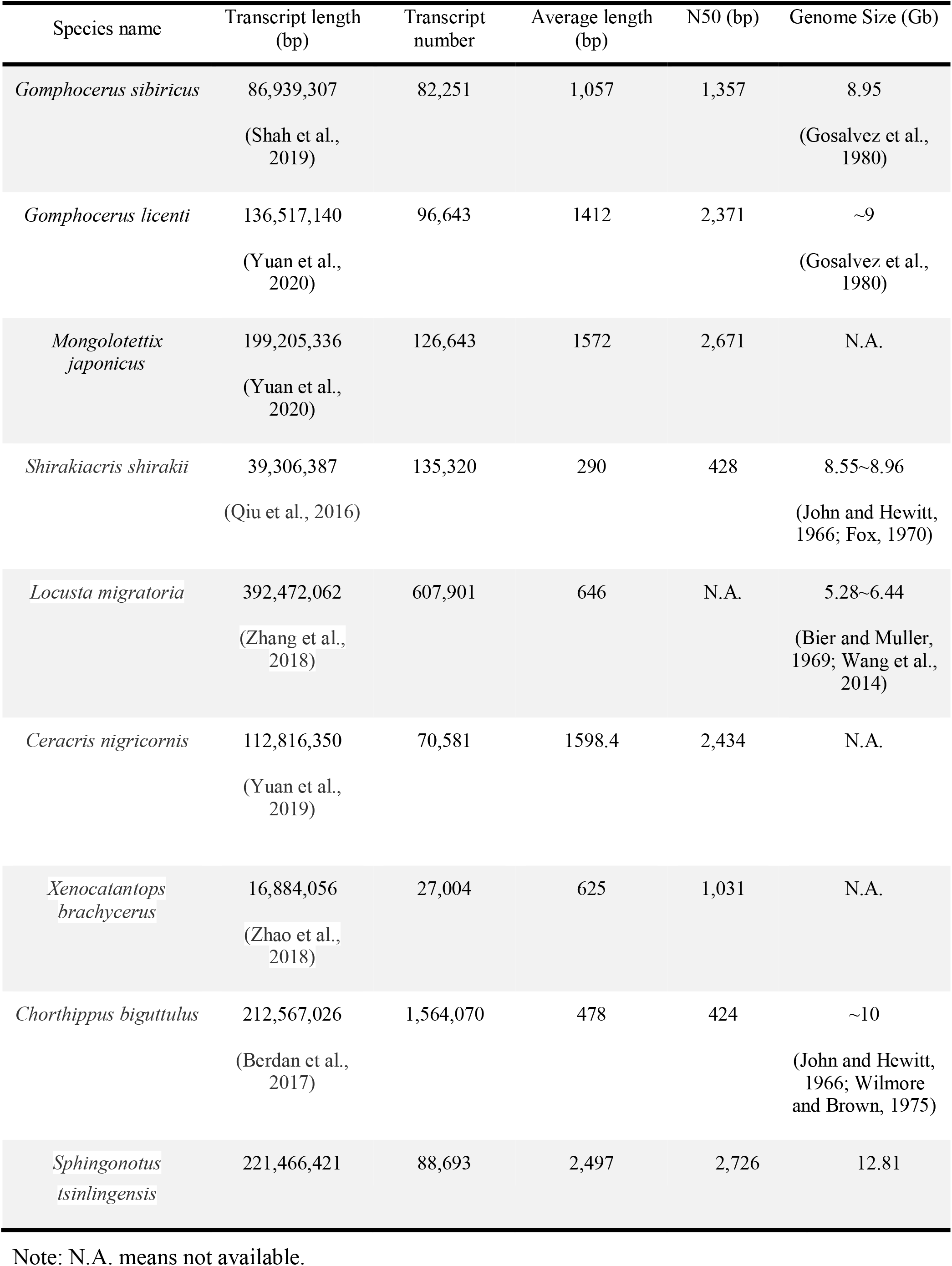
Comparison of transcriptome assemblies and gene numbers with the eight published Acrididae transcriptomes.

### 3.3 Prediction of coding sequences

From the full-length, non-chimeric isoforms of *S. tsinlingensis,* 48,502 transcripts with CDSs were identified, which accounted for ~54.68% of the total isoforms. The CDSs N50 length was 1,230 bp and each CDS encoded 229.3 amino-acids, on average (**Supplementary Figure S2**). A total of 37,569 CDSs were retrieved after integrated annotations from seven gene databases, accounting for 78.18% of the total CDSs (**Figure 2A**). A Venn diagram was created to show the contribution of different databases in the annotations; there were 3,575 genes commonly annotated by the five most commonly employed databases (**Figure 2B**).

**FIGURE 2.**
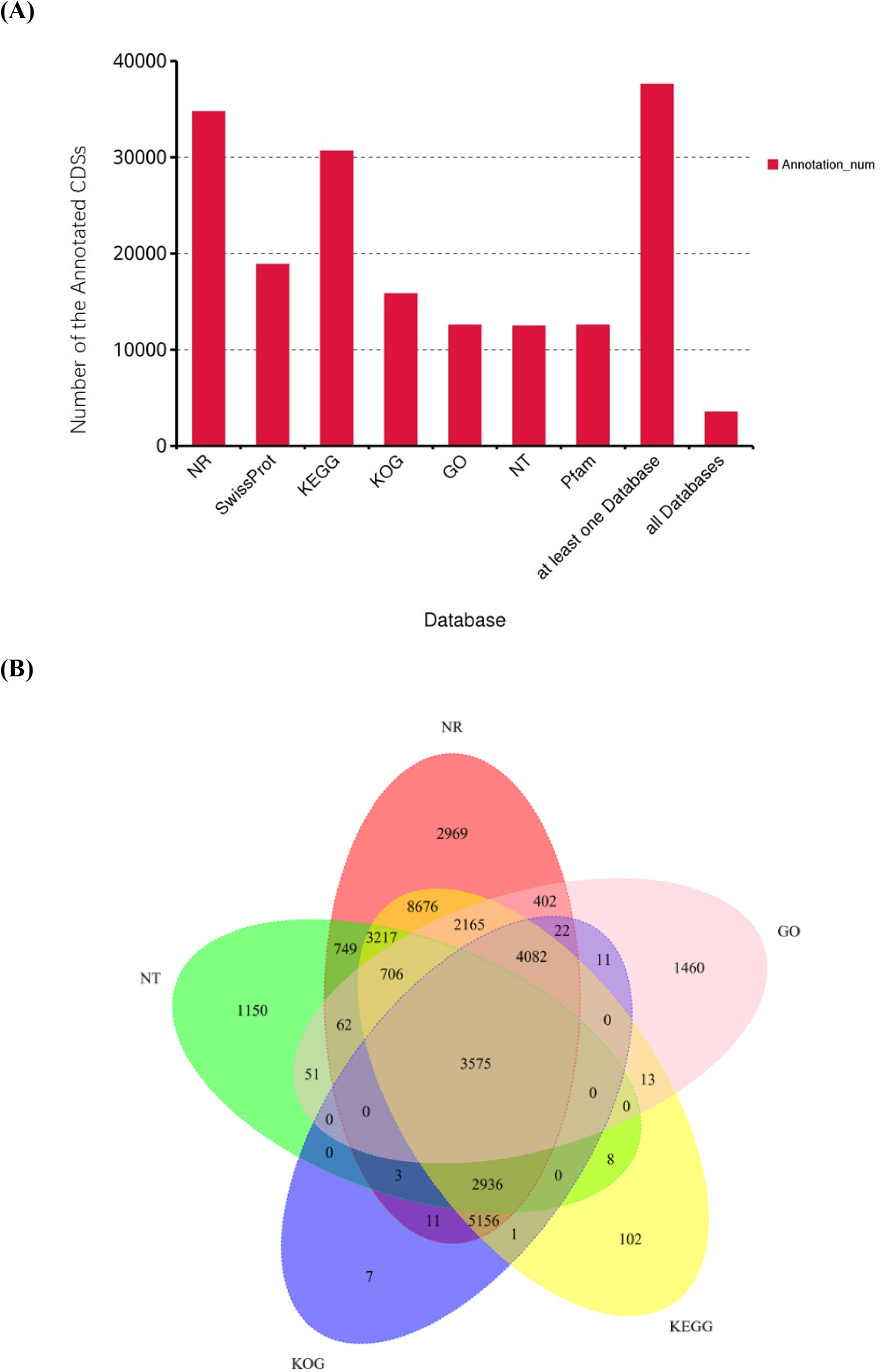
Statistics on the functional annotation of all *S. tsinlingensis* protein coding sequences (CDSs) in seven common databases (**A**), and the Venn diagram showing functional annotation of the *S. tsinlingensis* transcripts in the five most commonly used databases (**B**). NR, non-redundant protein sequence; KEGG, Kyoto Encyclopedia of Genes and Genomes; KOG, EuKaryotic Orthologous Groups; GO, gene ontology; NT, non-redundant nucleotide sequences; Pfam, protein family.

The main annotations were from the NR database (34,731 genes). Among them, *S. tsinlingensis* had most hits with *Zootermopsis nevadensis* (12.86%, 4459) (**Figure 3A**). The pattern of hits suggested that these species shared close phylogenetic positions, while the percentages corresponded to the number of their indexed sequences in the database, which also revealed that the proportion of *Acrididae* was not as high as that of other species. Gene ontology and KEGG annotations were adopted to describe the genome composition of *S. tsinlingensis.* **Cells** and **Cell Parts** (2,865 genes) were the most represented term in the cell component categories, while **Binding** (7,457 genes) was the most represented molecular function (**Figure 3B**). In the KEGG annotations, 30,637 annotations were assigned into 355 signaling pathways. The most enriched KEGG pathways were also related to basic life metabolism, including: **Signal Transduction** (1,008 genes), **Transport and Catabolism** (608 genes), **Amino Acid Metabolism** (606 genes), **Cancers Overview** (578 genes) and **Endocrine System** (571 genes) (**Figure 3C**). There were several highly conserved signaling pathways, critical in insect growth and body development, which were well enriched, including: Wnt (ko04310), Notch (ko04330), transforming growth factor-β (ko04350), Janus kinase/signal transducer and activator of transcription (ko04630), mitogen-activated protein kinase (ko04010), and Hedgehog (ko04340).

**FIGURE 3.**
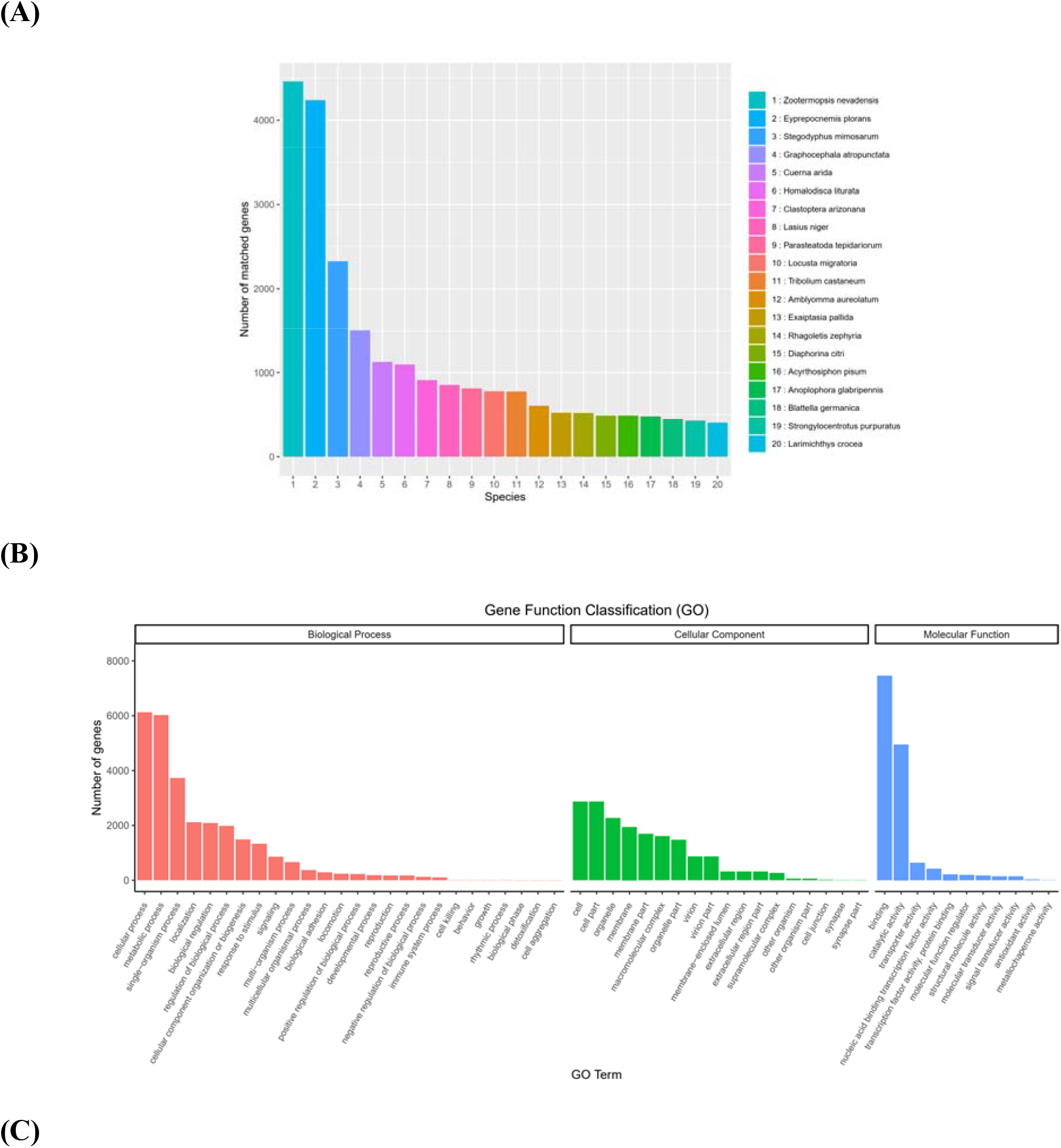
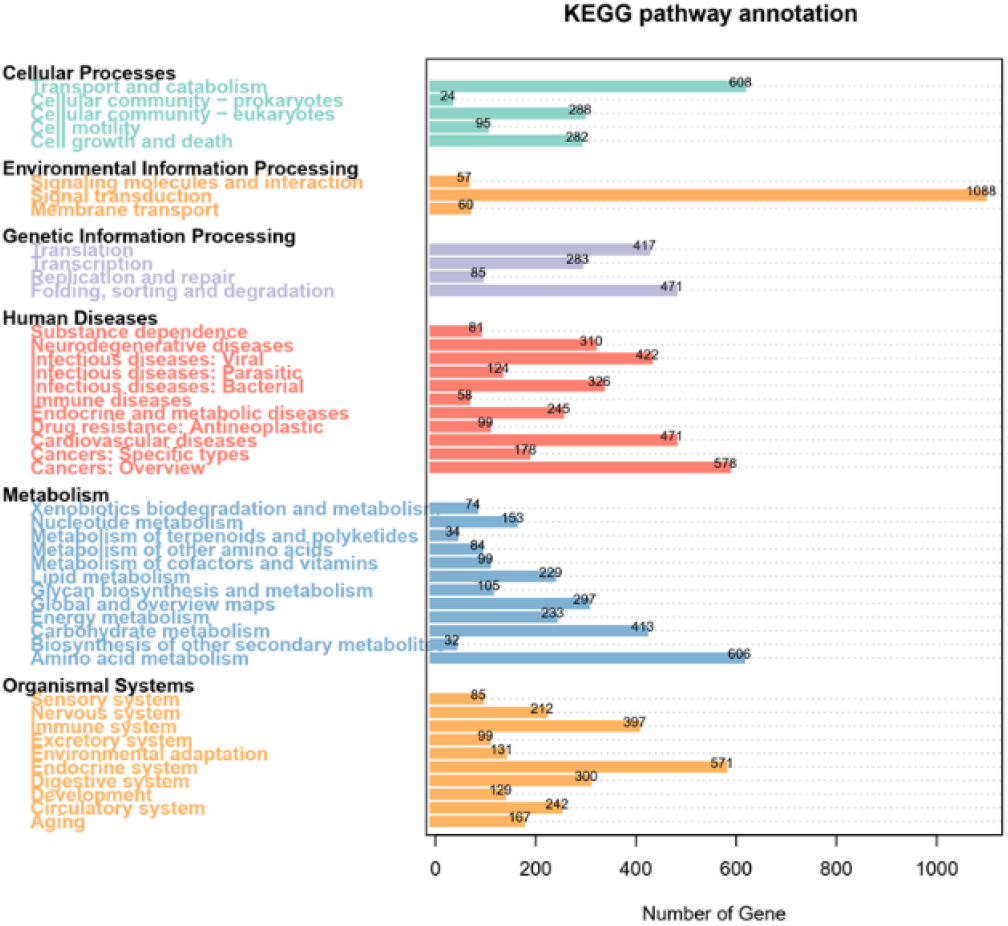
Functional annotation using the non-redundant protein sequence (NR) (**A**), gene ontology (GO) (**B**) and Kyoto Encyclopedia of Genes and Genomes (KEGG) databases (**C**).

### 3.4 Heat shock protein (HSP) genes and CYP450 genes

Drought-adaptive-related genes and their annotations were identified from the isoform sequences of *S. tsinlingensis,* including 70 HSP genes and 61 xenobiotic CYP450 genes. The gene length of HSP and CYP450 sequences varied between 1,438–4,428 bp, and 1,019–5,346 bp, respectively. Most BLAST identities for these target-focused sequences were below 95%, which suggested that they were novel to the current understanding of genetic mechanisms in *S. tsinlingensis* (**Figure 4**).

**FIGURE 4.**
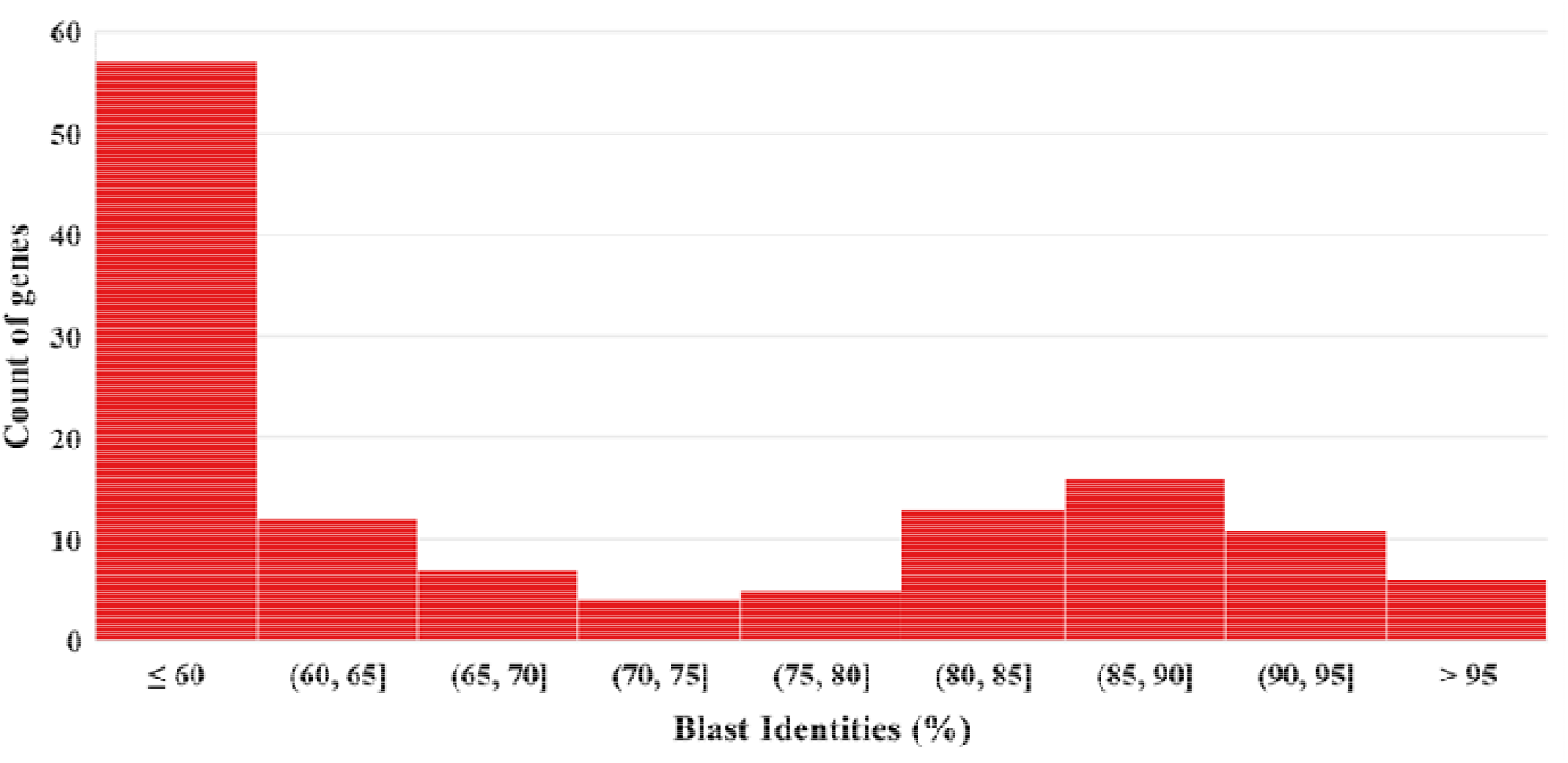
The basic local alignment search tool (BLAST) identities of heat shock protein (HSP) and CYP450 genes in the non-redundant protein sequence (NR) database.

### 3.5 LncRNA identification

LncRNAs were identified using a combination of the Coding Potential Calculator, Coding-NonCoding Index, Coding Potential Assessment Tool and Protein family (Pfam) methods. A total of 12,765 (22.94%) valid lncRNAs (with lengths > 200 bp and with more than two exons) were identified (**Figure 5**). A length distribution analysis of lncRNAs revealed that their lengths ranged from 212 −8,365 bp with a mean length of 2,090 bp. The N50 length of these identified lncRNAs was 2,185 bp.

**FIGURE 5.**
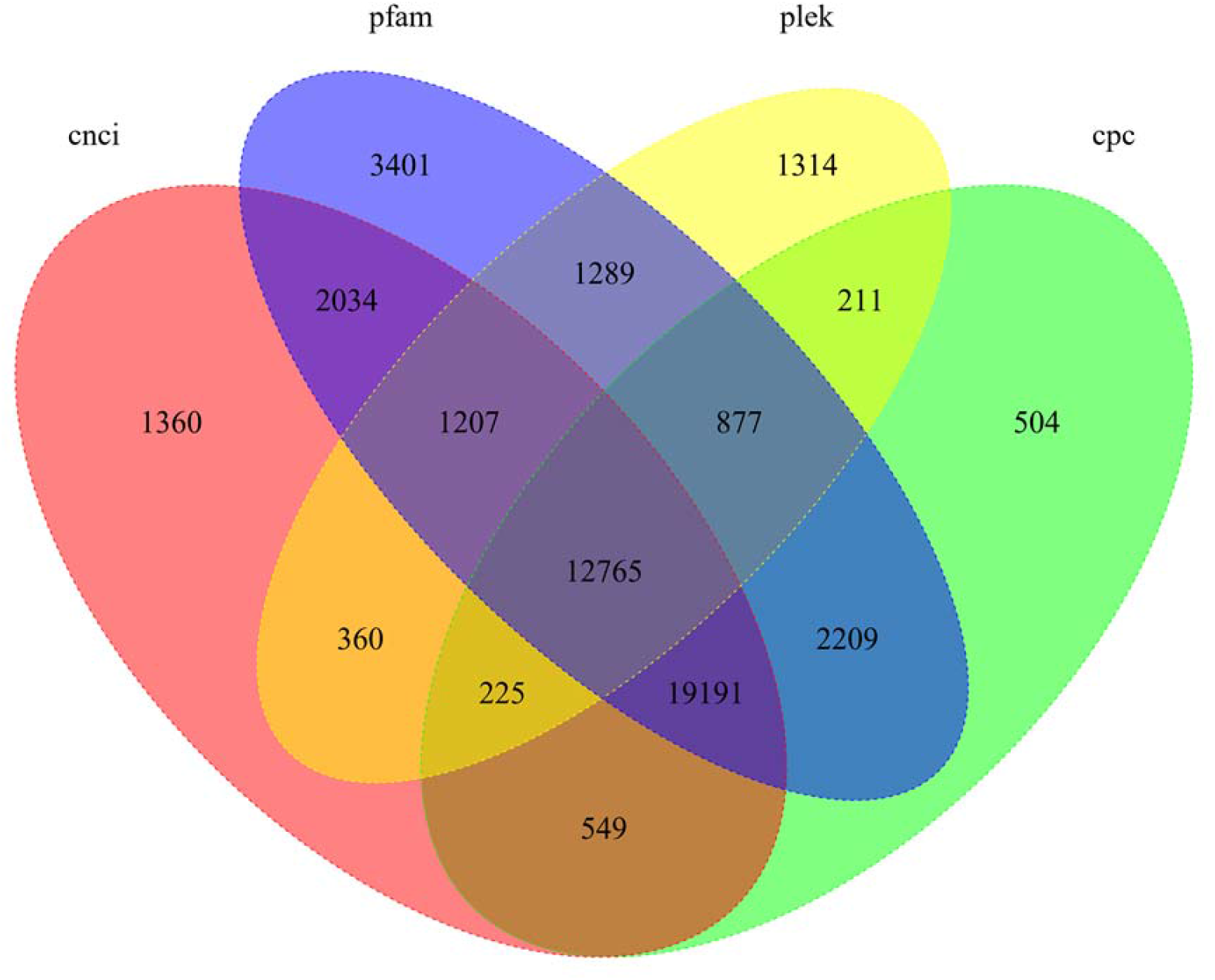
Venn diagram showing the distribution of long non-coding RNAs (lncRNAs) identified in *S. tsinlingensis.* CNCI, Coding-Non-Coding Index; Pfam, protein family; PLEK, ***p***redictor of ***l***ong non-coding RNAs and m***e***ssenger RNAs based on an improved ***k***-mer scheme; CPC, Coding Potential Calculator.

It was noticed that the number of lncRNAs was much fewer than that of mRNAs. To verify the two possible conditions: each kind of lncRNA corresponded to multiple mRNAs, or the valid lncRNA was not fully determined, the linkages between the 12,765 lncRNAs and 48,502 CDSs were checked. The lncRNAs only related to 9,347 CDSs, suggesting that the lncRNAs were not fully sequenced. Regarding the HSP and P450 genes, there were 4 and 0 related lncRNAs identified, respectively, suggesting that the regulation mechanism of these HSP and P450 genes was not well characterized.

### 3.6 TF identification

TFs participate in gene expression regulation by linking lncRNAs and mRNAs (Huang et al., 2018). Here, 414 putative TFs belonging to 41 TF gene families were predicted. The zf-C2H2 (20.29%, 84/414) was the most abundant TF family, followed by ZBTB (16.18%, 67/414) and THAP (15.22%, 63/414) (**Figure 6, Supplementary Table S3**). The relationships among the lncRNAs, TFs, and mRNAs were not discussed here due to the too few numbers of TFs. Instead, a gene ontology enrichment analysis was conducted using the genes where the TFs had been determined. Although smaller in numbers, these genes did not present functional bias and covered the basic metabolism similar to the whole genetic background. Survival-dependent mechanisms, such as multicellular organismal processes, developmental processes, and immune system processes were most significantly enriched (**Figure 7**).

**FIGURE 6.**
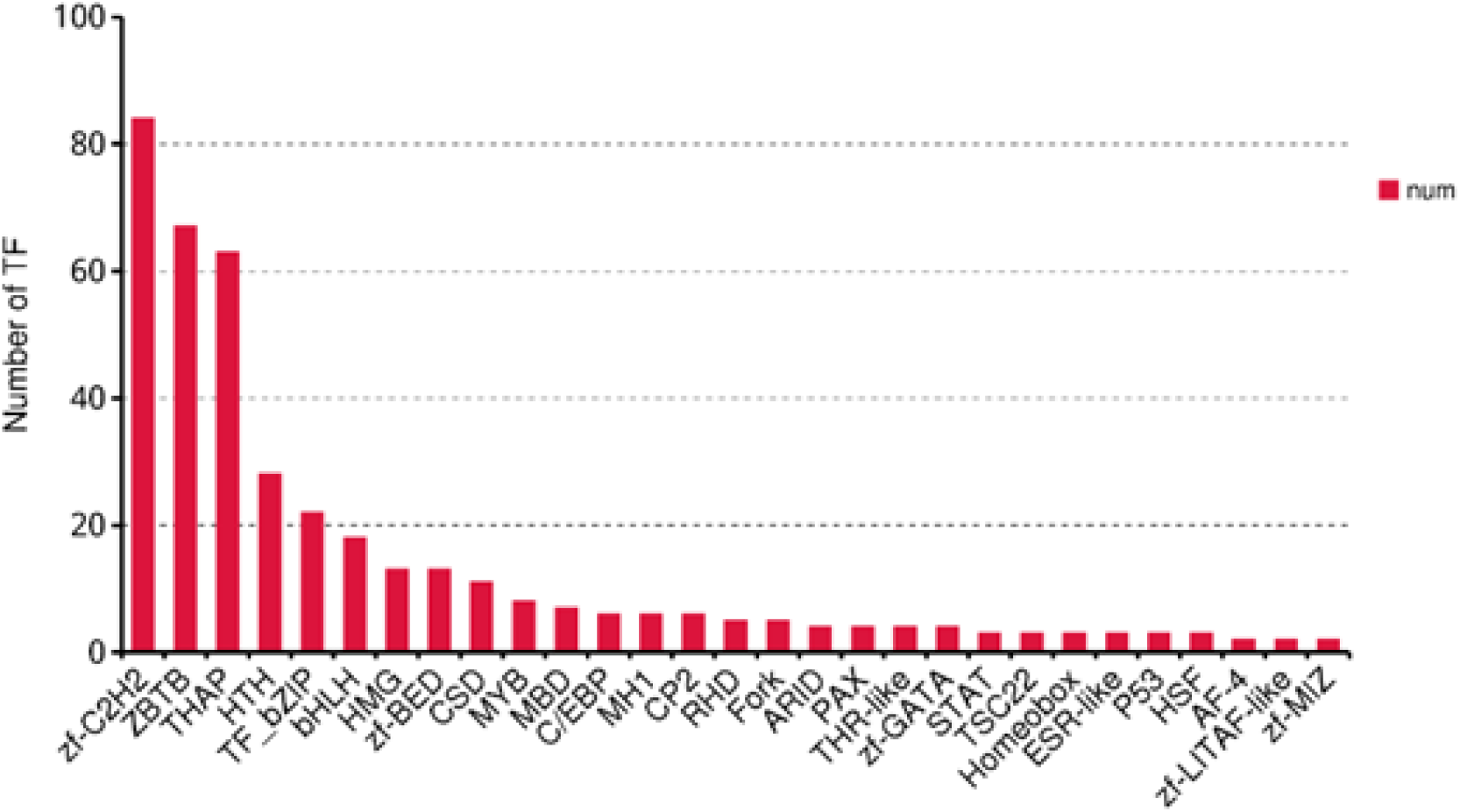
The types and number of the top transcription factors (TFs) identified in *S. tsinlingensis.*

**FIGURE 7.**
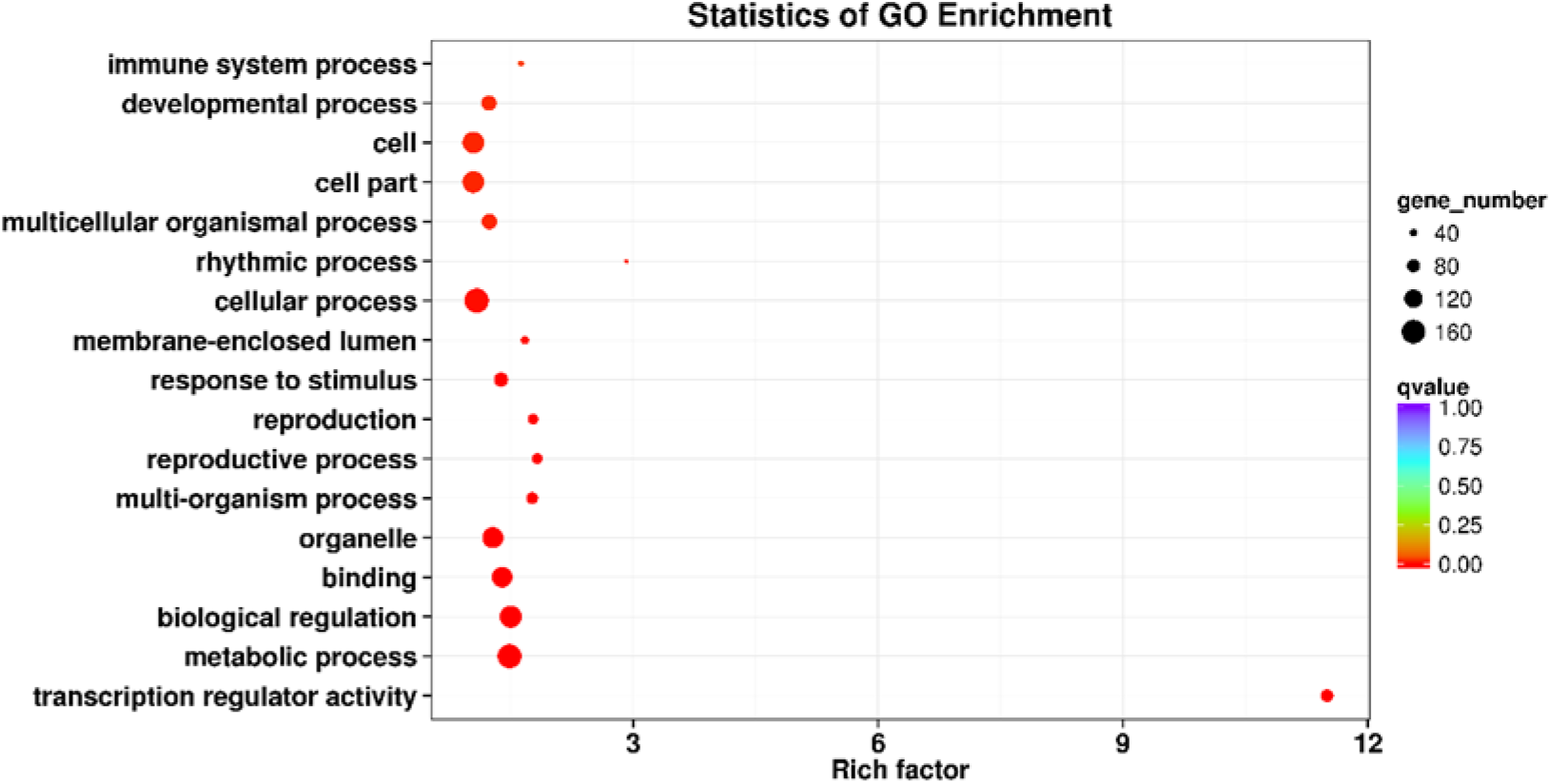
Gene ontology (GO) enrichment reveals the functions of genes that have been determined to be transcription factors (TFs). Note: Rich factor refers to the ratio of the number of differentially expressed transcripts to the total number of annotated transcripts located in the GO term, and the Qvalue is the Pvalue corrected by multiple hypothesis tests, with a value range of 0 to 1. The closer the Qvalue is to zero, the more significant the enrichment is.

### 3.7 SSR motif analysis

SSRs are important molecular markers that reflect genetic polymorphism (Xiao et al., 2015). A total of 36,488 SSRs were identified in all isoforms of *S. tsinlingensis.* A total of 3,719 genes contained more than one SSR, and 328 SSRs were present in a compound form. Among six SSR types, mononucleotide repeats (18,078, 49.55%) were the most abundant, followed by di-nucleotide repeats (9,859, 27.02%), tri-nucleotide repeats (7,612, 20.86%), tetra-nucleotide repeats (791, 2.17%), pentanucleotide repeats (107, 0.29%), and hexa-nucleotide repeats (41, 0.11%) (**Figure 8**). This full-length transcriptome of *S. tsinlingensis* contained more SSRs than the other known full-length transcriptomes of grasshoppers, including *G. licenti, M. japonicus,* and *S. shirakii,* possibly because of the higher heterozygosity of *S. tsinlingensis* or a more thorough SSR search.

**FIGURE 8.**
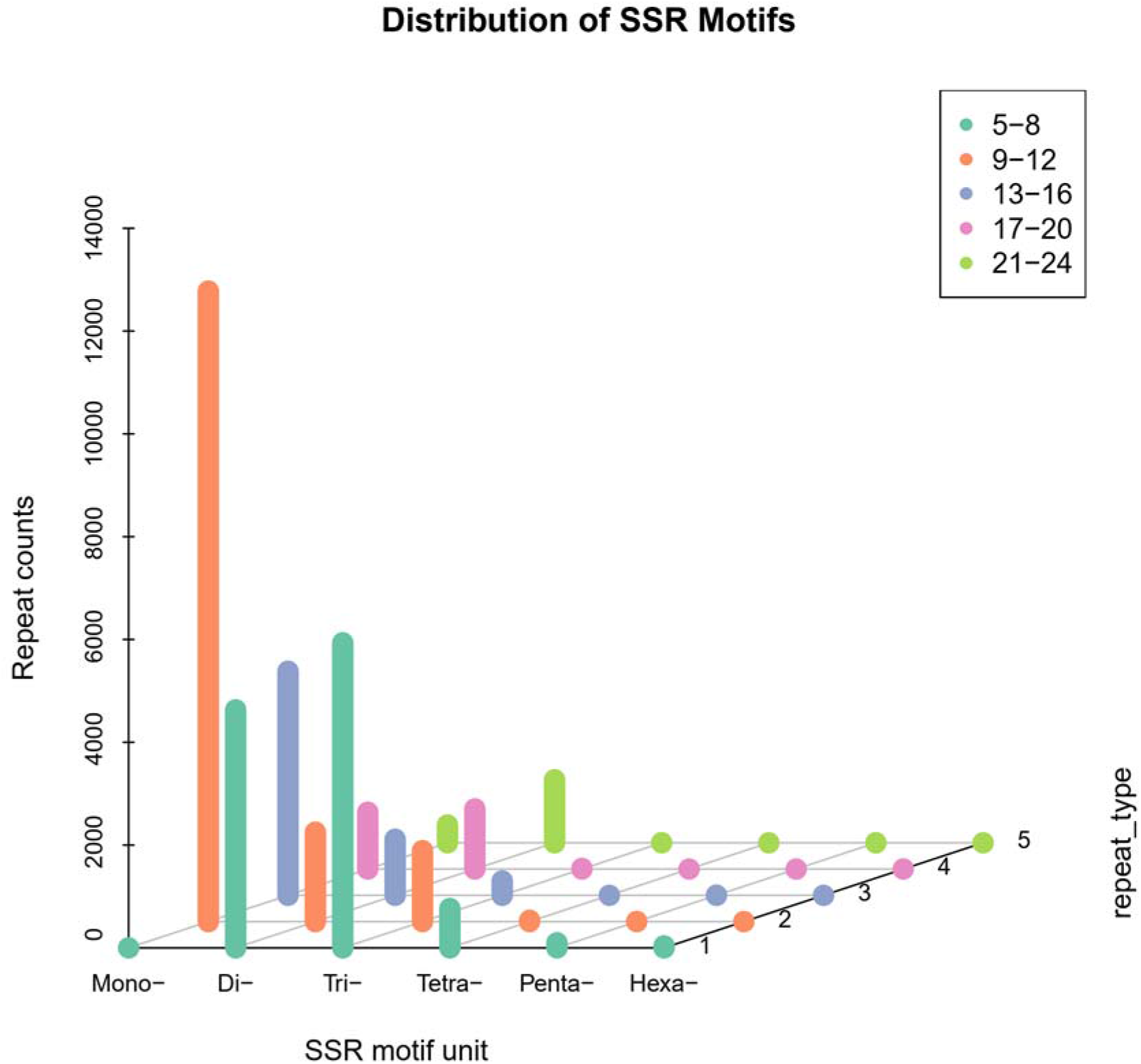
Distribution of simple sequence repeat (SSR) motifs in *S. tsinlingensis.* The X-coordinate represents the SSR type, the Y-coordinate represents the repeat type with a specific repetition number, and the Z-coordinate represents repeat counts.

## 4 Discussion

This GS and transcriptome of *S. tsinlingensis* is the first genetic study in *Sphingonotus spp.* Although *S. tsinlingensis* is an oriental species endemic to China, it represents the most popular morpholgical type of the grasshopper genus *Sphingonotus* and the subgenus *S. Spingonotus,* whose hind wing possesses a curved black bind and blue-color on the base (Zheng et al., 1963). This subgenus contains most species of the genus and is distributed in all regions of the genus (Husemann et al., 2014; Husemann et al., 2015). Therefore, the genome and transcriptome may provide a reference for the study of the evolution and ecological adaptation of the *Sphingonotus* species. These data, such as lncRNAs, TFs, and SSRs may be used as a genetic background, for gene identification, homologous gene screening, phylogenetics, and adaptive evolution analysis.

Based on the ultra-large GS, we suggest using the full-length transcriptome instead of the whole genome when conducting genetic studies on the grasshoppers of this genus. In this study, the full-length transcriptome was successfully obtained through double-depth sequencing. The success of this strategy will help to optimize the experimental design for future similar studies. Because grasshoppers have very large genomes, and genome assemblies are difficult to construct for highly repetitive regions (Schatz et al., 2010; Shah et al., 2019), grasshopper genome assembly requires a lot of sequencing and computation, and is time-consuming and expensive. Even when sequencing mRNA, the sequencing often fails to enrich effective sequences due to the large GS, and rare genes are missing due to sequencing preferences (Gao et al., 2016). To solve this problem, in this study, the depth of sequencing was increased to obtain enough effective sequences. The number of protein expressing genes in the functional gene dataset obtained in this study was the same as that in previous studies. The proportion of annotated functional genes increased, and the average length of genes was greatly improved. From the composition of gene functions, gene detection was complete, and many basic metabolism-related genes were detected. In particular, of the grasshopper species that reported both the transcriptome and genome, the GS of *S. tsinlingensis* was the largest. Its GS ~12.81 Gb is almost twice the size of *L. migratoria* (Wang et al., 2014). A comprehensive excavation of the effective sequences in the genome was conducted when the ratio of transcript size against GS was less than 2%.

Although these results indicated that the full-length transcriptome was an effective method to study the genomics of grasshopper species with ultra-large and complex genomes, some limitations were also discovered. For example, lncRNAs, miRNAs, TFs, and mRNAs obtained from the full-length transcriptome usually play important regulatory roles and construct mRNA-miRNA-lncRNA and TF interaction networks (Ye et al., 2018). However, in this study, an association among these four regulatory elements was not identified. The regulatory elements detected by functional enrichment only reflected the basic metabolic processes of the body. To obtain better data in future studies and to better enrich the results of the full-length transcriptome, a combination of multi-omics methods should be used, such as competing endogenous RNA for identification (Jiang et al., 2020), supplemented with mRNA next-generation sequencing for expression profile analysis.

This is the first GS and full-length transcriptome study on the grasshopper genus *Sphingonotus*. *S. tsinlingensis* had a 12.81 Gb ultra-large genome. Deep-depth Pacbio sequencing is the most optimized method to retrieve nucleic acid sequences from species with ultra-large and complex genomes. The full-length transcriptome in this study provides a reference resource for future studies of gene identification and comparison, and will help to improve the understanding of phylogeny, development, evolution, and adaptation mechanisms of grasshoppers.

## Supporting information

All supplementary materials

## 6 Author Contributions

LZ and D-LG conceived the study and designed the experiments. LZ, KS, LW, and HW performed the sequencing experiments. D-LG and LZ analyzed the data. LZ and D-LG wrote the manuscript. D-LG and S-QX revised the manuscript. All authors read and approved the final manuscript.

## 8 Acknowledgments

Professor Sheng-Quan Xu and Dr. De-Long Guan are co-corresponding authors.

## 10 Supplementary Material

Table S1. Summary of filtered reads.

Table S2. Statistics of functional annotations.

Table S3. Statistics of transcription factors.

Figure S1. Length distribution of clean-reads. A: subreads length distribution; B: circular consensus sequence (CCS) length distribution; C: full-length non-chimeric (FLNC) read length distribution; D: consensus read length distribution.

Figure S2. Length distribution of all CDS.

## 11 Data Availability Statement

The CCS sequences were uploaded in the NCBI SRA database with the accession number PRJNA707366. The FLNCs and extracted protein-coding sequences, along with their corresponding functional annotations were released to Zenodo server (Doi: 10.5281/zenodo.4588269).

