## Supplementary material for "Genome Size Estimation and Full-length Transcriptome of *Sphingonotus tsinlingensis:* Genetic Background for the Drought-Adapted Grasshopper": All supplementary materials

### Supplementary Tables

**Table S1.** Summary of filtered reads.

| Classification | Number | Average length (bp) | N50 (bp) |
| --- | --- | --- | --- |
| Polymerase Reads | 994,900 | 66,247 | 119,072 |
| Subreads | 28,612,043 | 2,229 | 2,410 |
| CCS | 901,383 | 2,556 | 2,770 |
| FLNC | 712,109 | 2,433 | 2,662 |
| Non-redundant isoforms | 88,693 | 2,497 | 2,726 |

CCS: circular consensus sequences; FLNC read: full-length non-chimeric read.

**Table S2.** Statistics of functional annotations.

| Database | NR | SwissProt | KEGG | KOG | GO | NT | Pfam |
| --- | --- | --- | --- | --- | --- | --- | --- |
| Annotation  number | 34,731 | 18,866 | 30,637 | 15,804 | 12,549 | 12,457 | 12,549 |

**Table S3.** Statistics of transcription factors.

| Transcription factors | Number |
| --- | --- |
| zf-C2H2; ZBTB; THAP; HTH; TF_bZIP; bHLH; HMG; zf-BED; CSD; MYB; MBD; C/EBP; MH1; CP2; RHD; Fork; ARID; PAX; THR-like; zf-GATA; STAT; TSC22; Homeobox; ESR-like; P53; HSF; AF-4; zf-LITAF-like; zf-MIZ | 84; (67; 63; 28; 22; 18; 13; 13; 11; 8; 7; 6; 6; 6; 5; 5; 4; 4; 4; 4; 3; 3; 3; 3; 3; 3; 2; 2; 2) |

### Supplementary Figures

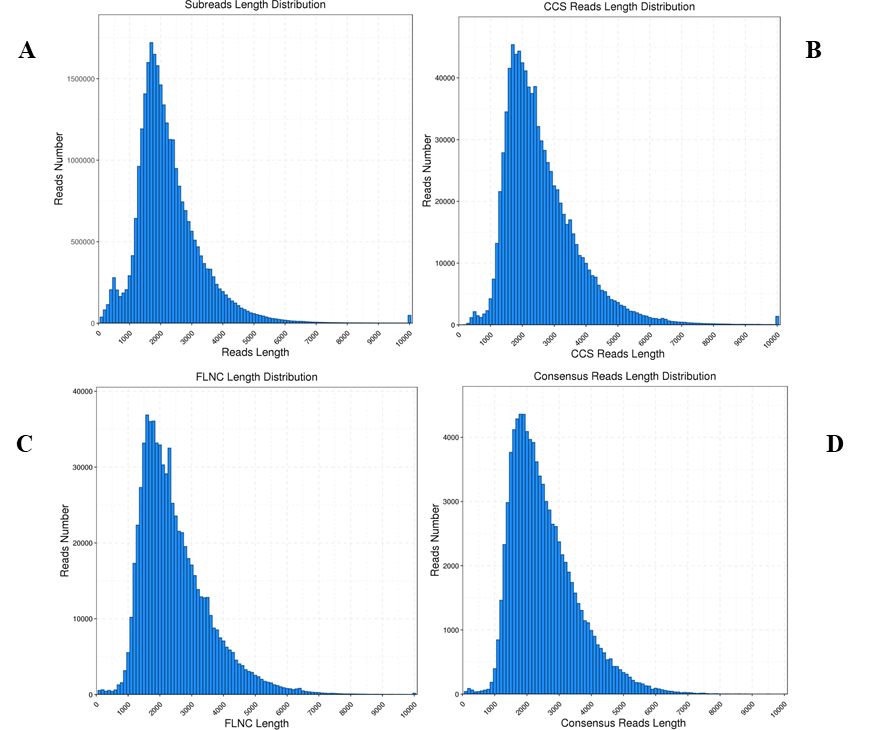

**Figure S1**. Length distribution of clean-reads. A: subreads length distribution; B: circular consensus sequence (CCS) length distribution; C: full-length non-chimeric (FLNC) read length distribution; D: consensus read length distribution.

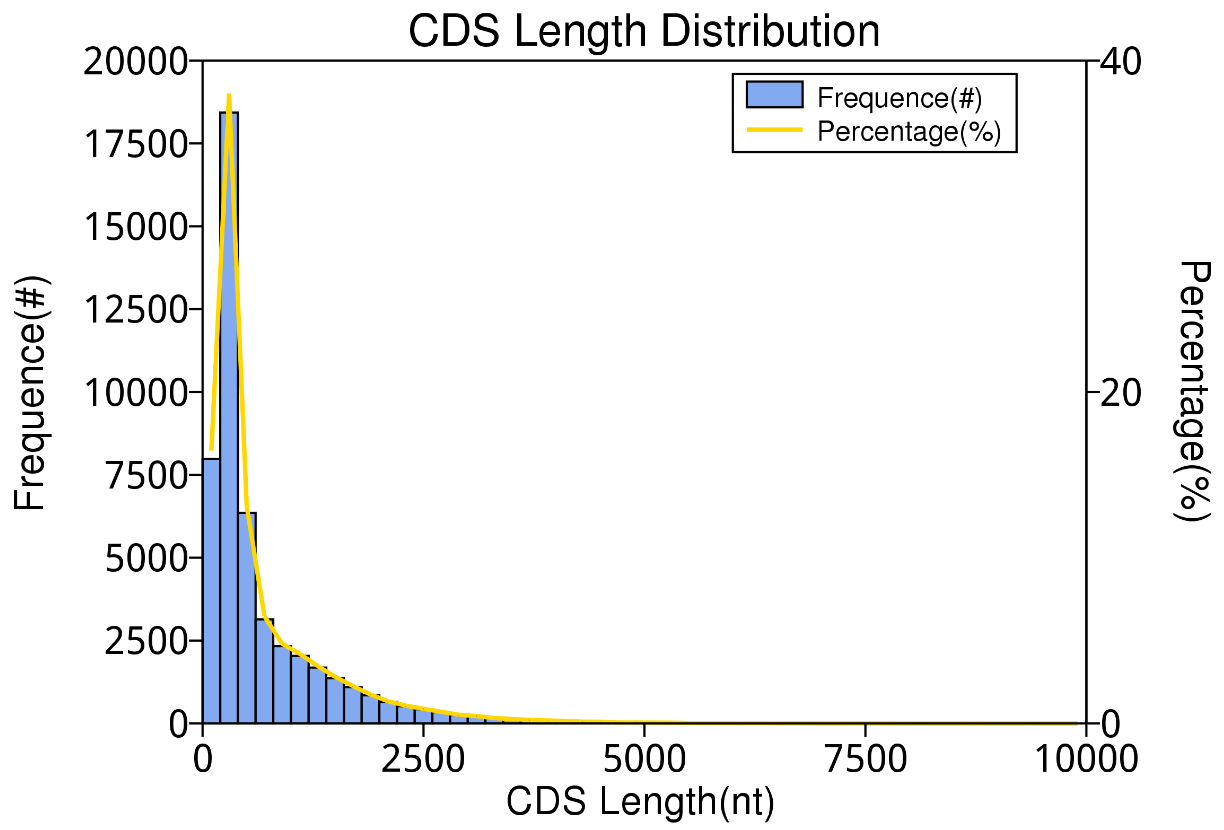

**Figure S2**. Length distribution of all CDS.
